# Fast evolution of SARS-CoV-2 BA.2.86 to JN.1 under heavy immune pressure

**DOI:** 10.1101/2023.11.13.566860

**Authors:** Sijie Yang, Yuanling Yu, Yanli Xu, Fanchong Jian, Weiliang Song, Ayijiang Yisimayi, Peng Wang, Jing Wang, Jingyi Liu, Lingling Yu, Xiao Niu, Jing Wang, Yao Wang, Fei Shao, Ronghua Jin, Youchun Wang, Yunlong Cao

## Abstract

While the BA.2.86 variant demonstrated significant antigenic drift and enhanced ACE2 binding affinity, its ability to evade humoral immunity was relatively moderate compared to dominant strains like EG.5 and HK.3. However, the emergence of a new subvariant, JN.1 (BA.2.86.1.1), which possesses an additional spike mutation, L455S, compared to BA.2.86, showed a markedly increased prevalence in Europe and North America, especially in France. Here, we found that L455S of JN.1 significantly enhances immune evasion capabilities at the expense of reduced ACE2 binding affinity. This mutation enables JN.1 to effectively evade Class 1 neutralizing antibodies, offsetting BA.2.86’s susceptibility and thus allowing it to outcompete both its precursor BA.2.86 and the prevailing variants HV.1 (XBB.1.5+L452R+F456L) and JD.1.1 (XBB.1.5+L455F+F456L+A475V) in terms of humoral immune evasion. The rapid evolution from BA.2.86 to JN.1, similar to the earlier transition from BA.2.75 to CH.1.1, highlights the importance of closely monitoring strains with high ACE2 binding affinity and distinct antigenicity, despite their temporarily unremarkable immune evasion capabilities. Such strains could survive and transmit at low levels, since their large antigenic distance to dominant strains allow them to target distinct populations and accumulate immune-evasive mutations rapidly, often at the cost of receptor binding affinity.

---

The SARS-CoV-2 saltation variant BA.2.86, which was quickly designated as a variant under monitoring (VUM) after its emergence, has garnered global attention. Although BA.2.86 did not exhibit significant humoral immune escape and growth advantage compared to current dominant variants, such as EG.5.1 and HK.3, it demonstrated remarkably high ACE2 binding affinity ^1-5^. This increased binding affinity, coupled with its distinct antigenicity, may enable BA.2.86 to accumulate immune-evasive mutations during low-level populational transmission, akin to the previous evolution from BA.2.75 to CH.1.1 and XBB ^6-9^. Notably, with just one additional receptor binding domain (RBD) mutation, L455S, compared to its predecessor BA.2.86, the JN.1 variant rapidly became predominant in France (Figure 1A and Figure S1), surpassing both BA.2.86 and the “FLip” (L455F+F456L) strains. A thorough investigation into the immune evasion capability of JN.1, particularly given its few additional mutations, is imperative.

**Figure.**
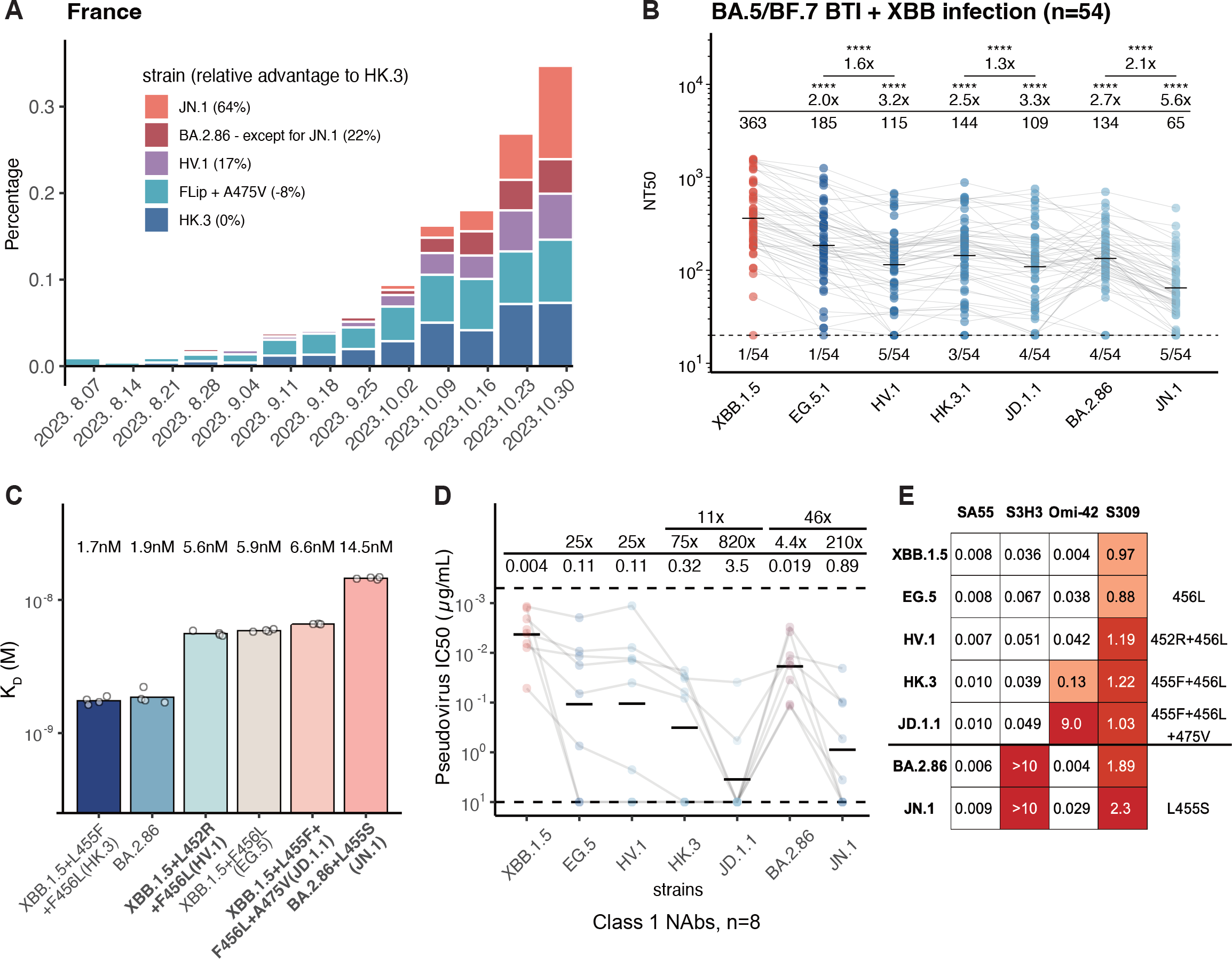
JN.1 exhibits profound immune evasion and decreased ACE2 binding affinity (A) Sequence percentages of prevalent variants since August 2023 including JN.1, BA.2.86 (the original BA.2.86 and its subvariants except JN.1), HV.1, FLip + A475V and HK.3. The growth advantages relative to HK.3 in past 2 month of these strains are denoted in the legend within parentheses. Data are collected from covSPECTRUM. (B) The 50% neutralizing titer (NT50) of convalescent plasma against SARS-CoV-2 variants measured in individuals who received three CoronaVac doses and had breakthrough infection with BA.5 or BF.7 followed by XBB reinfection (n = 54). Labels for geometric mean titers (GMT) are located above each group, with the fold changes and statistical significances indicated above the GMT labels. Below the dashed line are labels specifying the numbers of negative samples which are related to the limit of detection (NT50=20). Two-tailed Wilcoxon signed-rank tests of paired samples were used. *p<0.05, **p<0.01, ***p<0.001, and ****p<0.0001. (C) The human ACE2 (angiotensin-converting enzyme 2) binding affinities of HK.3 (XBB.1.5+L455F+F456L), BA.2.86, HV.1 (XBB.1.5+L452R+F456L), EG.5 (XBB.1.5+F456L), JD.1.1 (XBB.1.5+L455F+F456L+A475V), and JN.1 (BA.2.86+L455S) RBD determined by SPR sensorgrams. KD values (nM) are displayed above the bars, and all replicates are represented as points. (D) Class 1 neutralizing antibodies (NAbs) resistance against pseudovirus of XBB.1.5, EG.5, HV.1, HK.3, JD.1.1, BA.2.86, and JN.1 strains indicated by the IC50 values (n=8). The values and related fold changes when compared to D614G or other strains are labelled. (E) The IC50 (ug/mL) of approved or candidate monoclonal neutralising antibody drugs targeting spike are assessed against XBB.1.5, EG.5, HV.1, HK.3, JD.1.1, BA.2.86, and JN.1 pseudovirus.

We first study the humoral immune evasion of JN.1 using pseudovirus-based neutralization assays with plasma from convalescent individuals from XBB infection. These individuals, having received three doses of inactivated vaccines, subsequently experienced XBB (XBB subvariants with S486P substitution) breakthrough infections (BTIs). Our study included two cohorts, one with 27 participants who had post-vaccination XBB BTI and another with patients reinfected with XBB after BA.5 or BF.7 BTI (Table S1). Importantly, JN.1 displayed significantly enhanced immune escape compared to BA.2.86 (Figure 1B). This was evidenced by a 2.1-fold decrease in 50% neutralization titers (NT50) among XBB reinfected post-BA.5/BF.7 infection individuals and a 1.1-fold decrease in NT50 in the XBB BTI convalescents (Figure 1B and Figure S2). Additionally, JN.1’s plasma evasion surpassed that of competitive variants HV.1 (EG.5+L452R) and JD.1.1 (FLip+A475V). Notably, HV.1 and JD.1.1 also exhibited significantly lower susceptibility to convalescent plasma compared to their parental strains after acquiring L452R and A475V, respectively, explaining their growth advantages.

Subsequently, we measured the binding affinity between human ACE2 (hACE2) and the RBD of JN.1 using surface plasmon resonance (SPR) assays. A notable reduction in ACE2 binding affinity was observed in JN.1, indicating that its enhanced immune escape capabilities come at the expense of reduced ACE2 binding (Figure 1C). A475V mutation in JD.1.1 (XBB.1.5 + FLip + A475V) also slightly dampens binding affinity while enhancing immune evasion compared to HK.3 (XBB.1.5 + FLip). In contrast, L452R of HV.1 did not significantly affect receptor-binding affinity.

Considering that the L455 is predominantly located at the epitope of RBD Class 1 antibodies, as indicated by prior research, we further examined the evasion capabilities of JN.1 in response to eight XBB.1.5-neutralizing Class 1 monoclonal antibodies (mAbs) ^7^. Pseudovirus neutralization assays demonstrated that the addition of the L455S mutation notably enhanced JN.1’s ability to evade Class 1 antibodies (Figure 1D). This mutation effectively compensated for BA.2.86’s susceptibility to this antibody group (Figure 1D). Similarly, FLip+A475V (JD.1.1) also exhibited increased resistance to Class 1 antibodies compared to the FLip variant (HK.3), offering insights into the recent trend of convergent A475V mutations among FLip variants. In terms of therapeutic antibodies, SA55 retained its neutralizing efficacy against all examined variants including JN.1 (Figure 1E). Together, these findings suggest that L455S greatly enhanced JN.1’s resistance to humoral immunity through significant compensation of BA.2.86’s weakness to Class 1 antibodies.

To sum up, JN.1, by inheriting BA.2.86’s antigenic diversity and fast gaining of L455S, rapidly achieved extensive resistance across RBD Class1/2/3 antibodies ^1^, and demonstrated higher immune evasion compared to BA.2.86 and other evasive strains like HV.1 and JD.1.1, at the expense of reduced hACE2 binding. This evolutionary pattern, similar to the previous transition from BA.2.75 to CH.1.1 and XBB ^2,3,9^, highlights the importance of closely monitoring strains with high ACE2 binding affinity and distinct antigenicity, like BA.2.86 and BA.2.75, despite their temporarily unremarkable immune evasion capabilities. Such strains could survive and transmit at low levels since their large antigenic distance to dominant strains would allow them to target distinct populations and have the potential to quickly accumulate highly immune-evasive mutations, often at the cost of receptor binding affinity.

## Acknowledgments

We are grateful to scientists in the community for their continuous tracking of SARS-CoV-2 variants and helpful discussion. We thank all volunteers for providing blood samples. This project is financially supported by the Ministry of Science and Technology of China (2023YFC3041500; 2023YFC3043200), Changping Laboratory (2021A0201; 2021D0102), and the National Natural Science Foundation of China (32222030, 2023011477).

## Author Contributions

Y.C. designed and supervised the study. S.Y. and Y.C. wrote the manuscript with inputs from all authors. Y.X. and R.J. recruited the SARS-CoV-2 convalescents. S.Y., F.J., W.S. and J.L performed sequence analysis and illustration. Y.Y. and Youchun W. constructed pseudoviruses. P.W., L.Y., Yao W., J.W. (BIOPIC), J.W. (Changping Laboratory) and F.S. processed the plasma samples and performed the pseudovirus neutralization assays. F.J., W.S., A.Y., X.N., and Y.C. analyzed the neutralization data.

## Methods

### Patient recruitment and plasma isolation

As previously described, blood samples from convalescent patients who had recovered from SARS-CoV-2 Omicron BTI or reinfection were obtained following approved study protocols (Ethics Committee archiving No. LL-2021-024-02 and No. 2022N045KY)^11,12^. Patients in the reinfection cohorts experienced first infections in December 2022 in Beijing and Tianjin and the second infection between May and June 2023^13^. The time of infections for individuals in the XBB BTI were between May and June 2023. The infections were confirmed by PCR test or antigen test while the viral strains of them were determined by sequencing.

After collection, whole blood samples were diluted 1:1 with PBS+2% FBS and then gradiently centrifugated with Ficoll (Cytiva, 17-1440-03). These plasma samples were then collected, aliquoted, stored at temperatures of −20 °C or lower, and subjected to heat inactivation before other experimental procedures.

### Pseudovirus neutralization assay

Based on the vesicular stomatitis virus (VSV) pseudovirus packaging system, pseudovirus of SARS-CoV-2 variant spike was generated^14^. G* Δ G-VSV virus (VSV G pseudotyped virus, Kerafast) was added to the cell culture supernatant., The the pcDNA3.1 vector incorporating spike gene which was optimized using a mammalian codon was transfected into 293T cells (American Type Culture Collection [ATCC], CRL-3216). After culture, the pseudovirus in the supernatant was harvested, filtered, aliquoted, and frozen at −80°C for subsequent use. Plasma samples or antibodies were serially diluted in culture media, then were with pseudovirus followed by a 1-hour incubation at 37 ° C in a 5% CO_2_ incubator. Digested Huh-7 cells (Japanese Collection of Research Bioresources [JCRB], 0403) were introduced into the antibody-virus mixture. After a day of incubation, the supernatant was removed. D-luciferin reagent (PerkinElmer, 6066769) was applied, left to incubate in darkness for 2 minutes. Then cells were lysed and transferred to the detection plates. A microplate spectrophotometer (PerkinElmer, HH3400) was used to detect the luminescence value and a four-parameter logistic regression model was employed to determine the IC50 values.

### Surface Plasmon Resonance

SPR measurements were performed on the constructed RBD of BA.2.86, JN.1 and XBB subvariants including HK.3,HV.1, EG.5 and JD.1.1 based on Biacore 8K (Cytiva). Human ACE2 tagged with Fc tag was immobilized onto Protein A sensor chips (Cytiva). After serial dilution (6.25, 12.5, 25, 50, and 100 nM), the purified RBD samples of SARS-CoV-2 variants were injected on sensor chips. Responses were captured by Biacore 8K Evaluation Software 3.0 (Cytiva) at room temperature, and the raw data were fitted to 1:1 binding model using Biacore 8K Evaluation Software 3.0 (Cytiva). Each strain underwent two or three independent replicates for validation.

## Supplementary Figures

**Figure S1.**
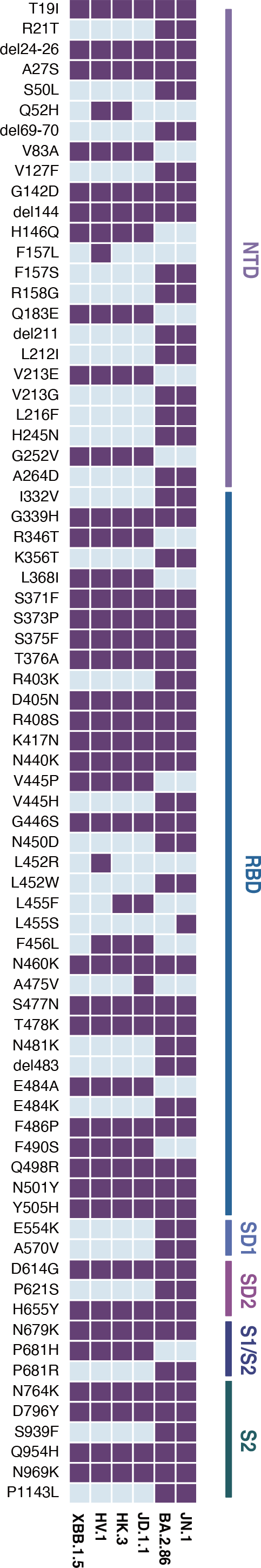
Sequence mutations of prevailing variants. Mutations of XBB.1.5, HV.1, HK.3, JD.1.1, BA.2.86 and JN.1 on the spike glycoprotein. The existence of mutations for each variant are indicated in purple. The sky-blue color denotes relatively absent mutations. The spike locations of listed mutations are labeled on the right.

**Figure S2.**
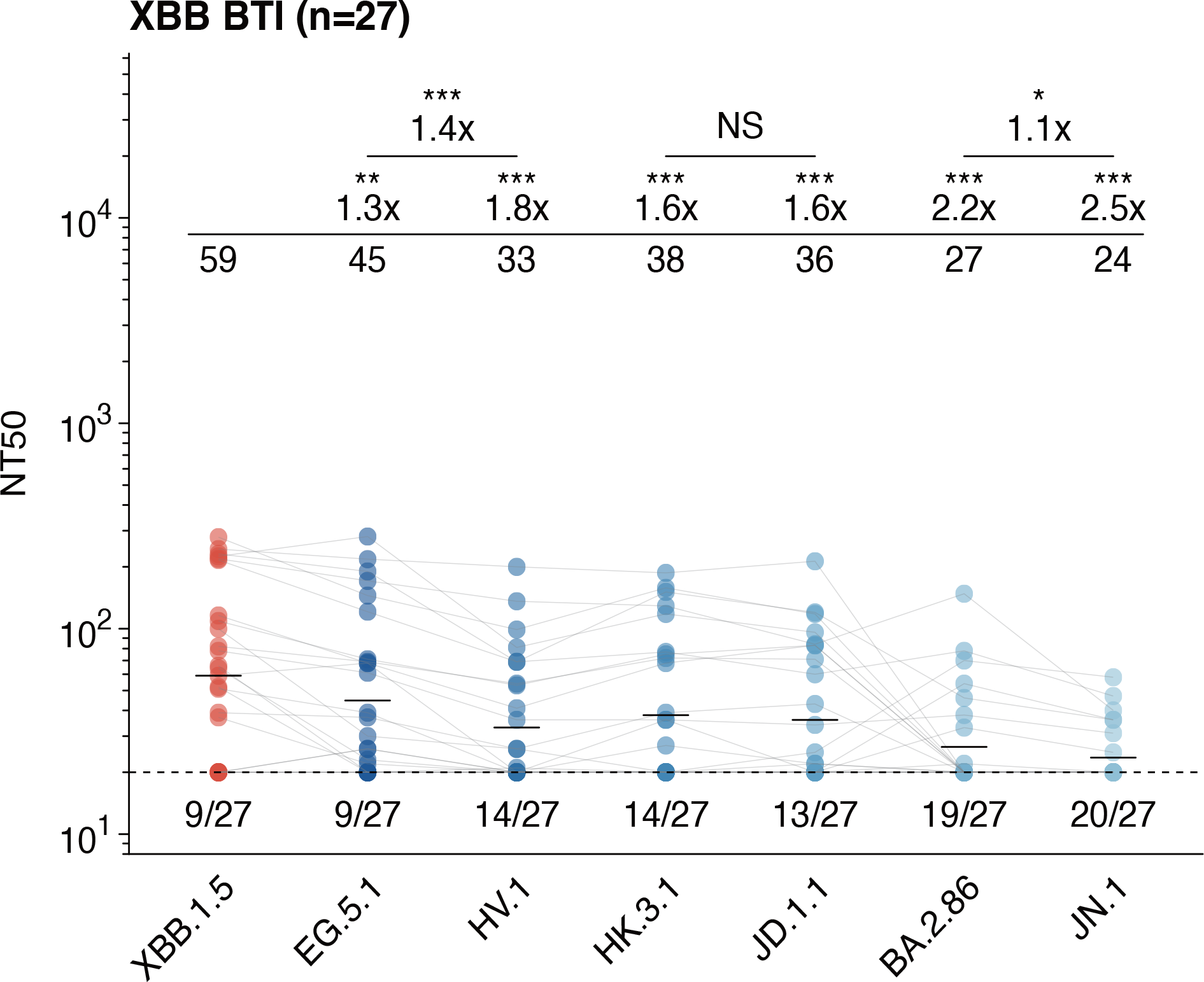
Neutralization of BA.2.86 against plasma from XBB BTI patients. Related to Figure B. NT50 against SARS-CoV-2 variants of convalescent plasma who received three CoronaVac doses and experienced XBB breakthrough infection (n = 27). The GMT values together with the relative fold changes and statistical significances are labelled above each group, and labels for the numbers of negative samples are below the dashed line which are related to the limit of detection (NT50=20). Two-tailed Wilcoxon signed-rank tests of paired samples were used. *p<0.05, **p<0.01, ***p<0.001, and ****p<0.0001.

## References

1. Yang, S. et al. Antigenicity and infectivity characterisation of SARS-CoV-2 BA.2.86. The Lancet Infectious Diseases 23, e457–e459 (2023).

2. Wannigama, D. L. et al. Tracing the new SARS-CoV-2 variant BA.2.86 in the community through wastewater surveillance in Bangkok, Thailand. The Lancet Infectious Diseases 23, e464–e466 (2023).

3. Wang, Q. et al. Antigenicity and receptor affinity of SARS-CoV-2 BA.2.86 spike. Nature 1–3 (2023) doi:10.1038/s41586-023-06750-w.

4. Uriu, K. et al. Transmissibility, infectivity, and immune evasion of the SARS-CoV-2 BA.2.86 variant. The Lancet Infectious Diseases 23, e460–e461 (2023).

5. Sheward, D. J. et al. Sensitivity of the SARS-CoV-2 BA.2.86 variant to prevailing neutralising antibody responses. The Lancet Infectious Diseases 23, e462–e463 (2023).

6. Cao, Y. et al. Characterization of the enhanced infectivity and antibody evasion of Omicron BA.2.75. Cell Host & Microbe 30, 1527-1539.e5 (2022).

7. Cao, Y. et al. Imprinted SARS-CoV-2 humoral immunity induces convergent Omicron RBD evolution. Nature 614, 521–529 (2023).

8. Yue, C. et al. ACE2 binding and antibody evasion in enhanced transmissibility of XBB.1.5. The Lancet Infectious Diseases 23, 278–280 (2023).

9. Wang, Q. et al. Evolving antibody evasion and receptor affinity of the Omicron BA.2.75 sublineage of SARS-CoV-2. iScience 26, 108254 (2023).

10. Jian, F. et al. Convergent evolution of SARS-CoV-2 XBB lineages on receptor-binding domain 455-456 enhances antibody evasion and ACE2 binding. 2023.08.30.555211 Preprint at 10.1101/2023.08.30.555211 (2023).

## Methods References

13. Pan, Y., Wang, L., Feng, Z., Xu, H., Li F., Shen, Y., Zhang, D., Liu, W.J., Gao, G.F., and Wang Q. (2023). Characterisation of SARS-CoV-2 variants in Beijing during 2022: anepidemiological and phylogenetic analysis. The Lancet 401, 664–672. 10.1016/S0140-6736(23)00129-0.

14. Li, Q., Wu, J., Nie, J., Zhang, L., Hao, H., Liu, S., Zhao, C., Zhang, Q., Liu, H., Nie L., et al. (2020). The Impact of Mutations in SARS-CoV-2 Spike on Viral Infectivity and Antigenicity Cell 182, 1284–1294.e1289. 10.1016/j.cell.2020.07.012.

